# Motor Cortex Excitability Reflects the Subjective Value of Reward and Mediates its Effects on Incentive Motivated Performance

**DOI:** 10.1101/387332

**Authors:** Joseph Galaro, Pablo Celink, Vikram S. Chib

## Abstract

Performance-based incentives tend to increase an individual’s motivation, resulting in enhancements in behavioral output. While much work has focused on understanding how the brain’s reward circuitry influences incentive motivated performance, fewer studies have investigated how such reward representations act on the motor system. Here we measured motor cortical excitability with transcranial magnetic stimulation (TMS) while female and male human participants performed a motoric incentive motivation task for prospective monetary gains and losses. We found that individuals’ performance increased for increasing prospective gains and losses. While motor cortical excitability appeared insensitive to prospective loss, temporal features of motor cortical excitability for prospective gains were modulated by an independent measure of an individual’s subjective preferences for incentive (i.e., loss aversion). Those individuals that were more loss averse had a greater motor cortical sensitivity to prospective gain, closer to movement onset. Critically, behavioral sensitivity to incentive and motor cortical sensitivity to prospective gains were both predicted by loss aversion. Furthermore, causal modeling indicated that motor cortical sensitivity to incentive mediated the relationship between subjective preferences for incentive and behavioral sensitivity to incentive. Together our findings suggest that motor cortical activity integrates information about the subjective value of reward to invigorate incentive motivated performance.

**SIGNIFICANCE STATEMENT:** Increasing incentives tend to increase motivation and effort. Using a motoric incentive motivation task and transcranial magnetic stimulation, we studied the motor cortical mechanisms responsible for incentive motivated motor performance. We provide experimental evidence that motor cortical sensitivity to incentive mediates the relationship between subjective preferences for incentive and incentive motivated performance. These results indicate that, rather than simply being a reflection of motor output, motor cortical physiology integrates information about reward value to motivate performance.

## INTRODUCTION

We modulate our performance according to the rewards at stake. Larger stakes tend to increase motivation, which in turn elicits increased behavioral output. Incentive motivation refers to the processes that convert higher reward expectancies into increased performance (Berridge, 2004). These processes include forming a subjective representation of prospective reward, which invigorates behavioral performance. The effects of incentive motivation on effortful exertion has been the topic of extensive investigation in psychology (Bolles, 1972; Bindra, 1974; Bolles and Fanselow, 1980), and, in more recent years, the field of cognitive neuroscience has begun to dissect how the brain’s reward circuity influences motivated performance (Pessiglione et al., 2007; Talmi et al., 2008; Chib et al., 2012; Schmidt et al., 2012). However, motivated performance is not only related to processing the rewards at stake, but also how these reward representations influence activity in motor cortex to result in behavioral performance. Despite the neural crosstalk between motivation and motor processing during incentivized performance (Mogenson et al., 1980; Bray et al., 2008; Talmi et al., 2008; Chib et al., 2014), the understanding of how motor cortical excitability gives rise to incentive motivated performance is fairly limited.

Transcranial magnetic stimulation (TMS) provides precise timing to study how motor cortical excitability is influenced by motivating stimuli. Freeman and colleagues recently used TMS to demonstrate that Pavlovian conditioned stimuli, paired with an instrumental response, served to increase motor cortical excitability and responding, while stimuli predicting the absence of reward did not invoke increases in motor excitability (Freeman et al., 2014). In a follow-up study, they found that presentation of aversive stimuli inhibited motor evoked potentials during trials that did not require instrumental responding (i.e. no-go trials) (Chiu et al., 2014). Together these results illustrate that motivational information spills into the motor system, influencing motor cortical excitability prior to execution.

Studies of binary choice have also used TMS to study the dynamics of motor excitability prior to action selection. This work has shown that motor cortical activity builds as a choice approaches and that excitability increases as function of the value of the chosen option (Duque and Ivry, 2009; Klein et al., 2012; Klein-Flügge et al., 2013). From these results it has been suggested that action selection during choice entails a competition, within motor-related areas, in which motor cortical excitability integrates reward value to drive a motor response. Furthermore, it was found that during binary choice of risky options, motor excitability was best described by chosen and unchosen subjective value (i.e., accounting for prospect theoretic measures) (Klein-Flügge and Bestmann, 2012). These studies suggest that the dynamics of motor excitability captures the value of reward during simple choice. However, it is not known how subjective-preferences for incentives might influence motor cortical excitability to drive incentive motivated instrumental responding.

The aim of this study was to investigate the role of motor cortical excitability on incentive motivation, and how these cortical processes interact with representations of subjective value to result in motivated performance. We hypothesized that the sensitivity of motor excitability to incentive would be predictive of an individual’s motivated performance. This hypothesis has its basis in previous TMS studies that found that motor cortical excitability, measured prior to instrumental responding, was modulated in response to conditioned stimuli that previously predicted appetitive and aversive outcomes (Chiu et al., 2014; Freeman et al., 2014; Freeman and Aron, 2016). We also hypothesized that motor cortical excitability would be related to an independent behavioral measure of subjective preferences for incentive. This hypothesis has its basis in previous TMS studies which found that motor cortical excitability reflected subjective chosen and unchosen values during binary choice (Klein-Flügge and Bestmann, 2012); and neuroimaging studies that found that the functional connectivity between reward regions and motor cortex, during instrumental responding for reward, was modulated by behavioral measures of subjective preferences (i.e., loss aversion) (Chib et al., 2012, 2014).

## MATERIALS AND METHODS

### Participants

All participants were right handed and prescreened to exclude those with a prior history of neurological or psychiatric illness. The Johns Hopkins Medical Institute Institutional Review Board approved this study, and all participants gave informed consent. Nineteen participants (mean age, 20; age range, 18–23; twelve females, seven males) were recruited and took part in the experiment. Each participant performed the motor task and a behavioral choice paradigm to characterize subjective preferences for incentive (i.e., loss aversion and risk aversion). One participant was excluded from the final analysis because of atypical choices during the behavioral choice paradigm (i.e. rejection of all gambles with potential losses).

### Experimental Setup and Brain Stimulation

Participants sat in a chair and held a force transducer (LMD300, Futek) between the thumb and forefinger of their right hand. During the experiment, participants rested their head in a custom-built gantry. The gantry minimized head-movements across trials and ensured accurate brain stimulator placement. An armrest ensured consistent positioning of the arm across trials. Visual stimuli were presented using MATLAB 2014a and Psychtoolbox-3 (Brainard, 1997; Kleiner et al., 2007).

To record motor evoked potentials elicited from TMS, surface electromyographic electrodes were placed on the first dorsal interosseous (FDI) muscle; signals were recorded, amplified, and filtered (Bortec Biomedical). To elicit motor evoked potentials, we delivered TMS using a 70mm figure-eight coil (Magstim) to the optimal scalp position over the left motor cortex. The coil was placed tangentially on the scalp with the handle pointing backward and laterally at a 45 angle away from the midline, perpendicular to the central sulcus. To ensure accurate and precise placement of the TMS coil throughout the experiment, we used a frameless neuronavigation system (Brainsight, Rouge Research) and coregistered participants’ heads to a default Talairach template provided in the Brainsight software suite. A deviation of more than 3mm or 15 degrees resulted in the experimenter repositioning the coil during the experiment. At the optimal motor cortex location, we determined the resting motor threshold, defined as the minimum TMS intensity that evoked a motor evoked potential (MEP) of 50 microvolts in 5 of 10 trials in the FDI of the right hand (Pascual-Leone et al., 1994; Rossini et al., 1994).

To control between participants and conditions, the stimulus intensity during testing was calibrated on a per subject basis such that the TMS pulse occurring 50ms following imperative cue on training trials elicited 1mV MEP. The stimulation intensity was fixed to this value for the remainder of the experiment (i.e., the same intensity was used on 50ms and 150ms trials in the later phases). This procedure was similar to those previously used to study pre-movement motor cortical excitability (Stefan et al., 2004; Vallence et al., 2013). To accomplish this, the first 30 trials of the unincentivized phase (described below) involved TMS pulses 50ms following imperative cue and the experimenter monitored the elicited MEPs to target 1mV. Additionally, the initial stimulator intensity was set to 120% of a participant’s resting motor threshold.

### Motor Task

Participants first performed a calibration phase to determine their maximum voluntary contraction (MVC) during an isometric pinch grip. This involved participants maintaining their maximum pinch exertion for 4 seconds, on 3 consecutive trials, each separated by a 5 second rest period. MVC was calculated as the maximum pinch force exerted on the 3 calibration trials. Since we acquired each individual’s MVC, we were able to standardize difficulty, based on MVC ability, across participants.

The main experiment was divided into two phases: unincentivized and incentivized (Figure 1). During both phases of the experiment participants performed an isometric pinch exertion task. This task was chosen because pinch grip isolates use of the FDI muscle, which we targeted in our TMS procedure, to study the relationship between incentives, motor excitability, and performance. TMS was performed on every trial of each phase of the experiment. Participants were instructed that they would receive a show-up fee of $15 dollars at the end of experiment in addition to any earnings from their performance in the incentivized phase.

**Figure 1.**
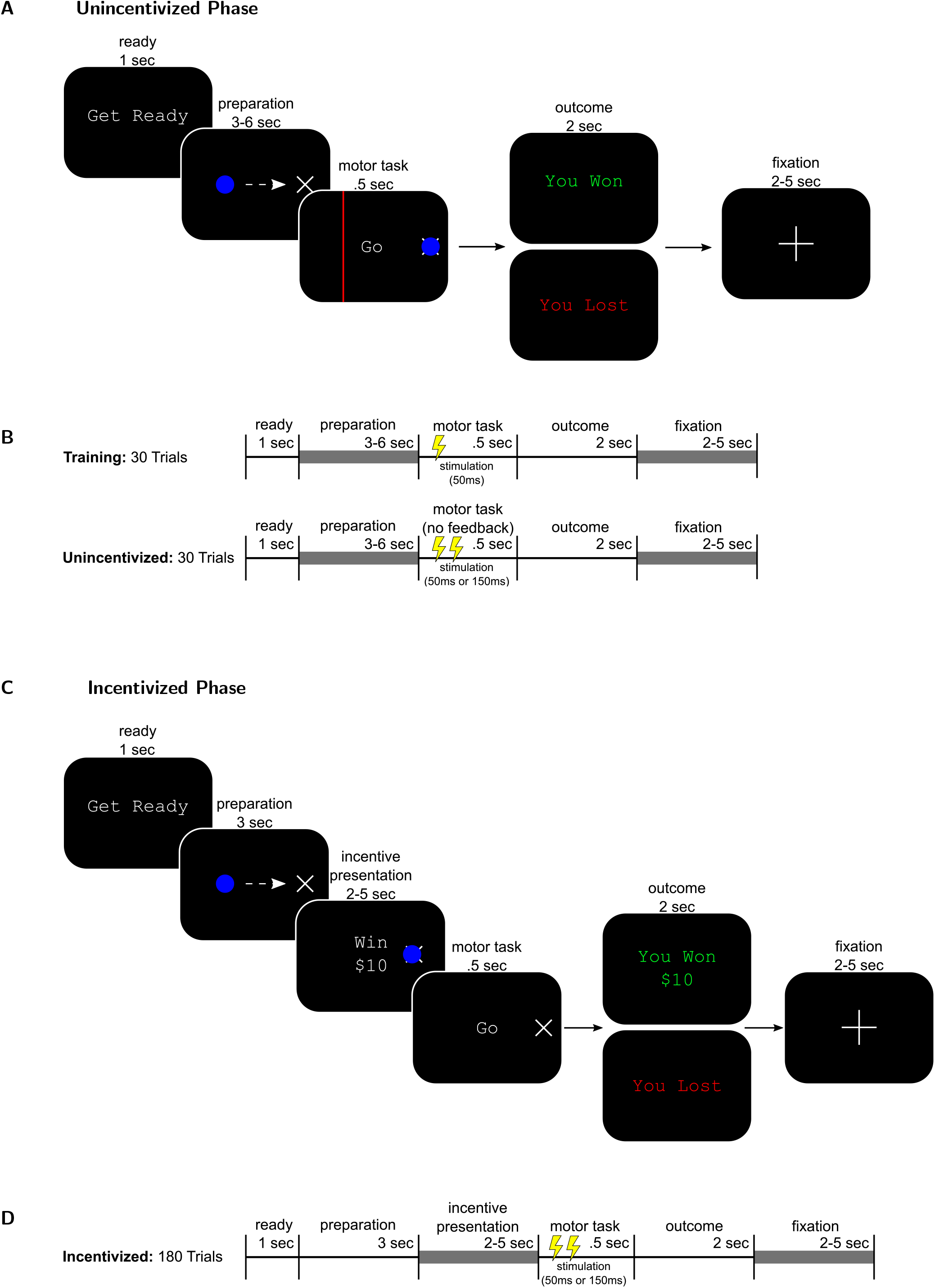
The incentive motivation motor task. **A**) Participants first performed an incentivized phase of the experiment to familiarize them with the requirements of behavioral paradigm and to calibrate TMS parameters. At the beginning of each trial, participants were presented a blue cursor that moved across the screen in proportion to the amount of pinch exertion. Squeezing the force transducer moved the cursor horizontally to the left, while relaxing caused the cursor to move to the right. To initiate the task, participants placed the cursor in the start position (×) for a random amount of time (3–6 s). The start position corresponded to minimal exertion while still grasping the transducer. During the task, a ‘Go’ cue and red target line appeared that was registered to 40% of MVC. To successfully achieve the task, participants had to move their cursor across the target line within 0.5 seconds. At the end of the trial they were shown a message indicating the outcome of their performance. In the case that a participant successfully moved the cursor across the target line, a positive message was displayed (“You Won”); otherwise, the participant was informed of her negative outcome (“You Lost”). **B**) The timeline of unincentivized trials. Participants first preformed 30 training trials in which stimulation occurred 50ms after the onset of the ‘Go’ Cue/motor task presentation. After training trials, stimulation was delivered for another 30 trials at either 50ms or 150ms after ‘Go’ cue/motor task presentation. **C**) Incentivized trials were identical to the unincentivized trials, except participants were presented with the incentive they were performing for, between the incentive presentation and motor task screens, and they were not given feedback of their cursor or the target line. **D**) The timeline of incentivized trials. There were a total of 180 incentivized trials and stimulation was delivered at either 50ms or 150ms after ‘Go’ cue/motor task presentation.

The unincentivized phase was comprised of 60 trials. At the beginning of each trial, participants were presented a blue cursor that moved across the screen in proportion to the amount of pinch exertion (Figure 1). Squeezing the force transducer moved the cursor horizontally to the left, while relaxing caused the cursor to move right. Participants were instructed to place the cursor in the start position (×) for a random amount of time (3–6 seconds). This start position corresponded to minimal pinch exertion while still grasping the force transducer. During the task, a ‘Go’ cue and a target line registered to 40% of MVC appeared on the screen. To successfully achieve the task, participants had to exert pinch effort to move the cursor across the target line within 0.5 seconds. At the end of a trial participants were shown a message indicating their performance. Following the initial 30 trial TMS calibration phase (described above), the remaining 30 trials involved TMS delivered at either 50ms or 150ms after presentation of the ‘Go’ cue, and visual feedback during exertion was withheld. The stimulation times were evenly distributed across trials.

During the incentivized phase participants performed the isometric pinch exertion task as described above, for varying amounts of monetary gain or loss. We did not present participants with feedback of their hand cursor or the effort target in order to allow them to reach the target effort level under their own implicit motivation. At the beginning of the experiment, participants were given an endowment of $20 in cash, separate from their show-up fee, and were told that at the end of the experiment, one trial would be selected randomly and a payment made according to their performance on that trial. Participants were told that their $20 endowment was given to them so that they could pay any eventual losses at the end of the experiment. This payout mechanism ensured that trials had significant monetary consequences and that participants evaluated each trial independently. Participants performed trials for a range of incentives (i.e. ± 0, 10, 20). Each incentive level was presented randomly 30 times for a total of 180 trials, with an equal balance of conditions for TMS pulse timing (i.e. 50ms TMS pulse; 150ms TMS pulse). At the beginning of each trial, participants were shown a message indicating the amount of incentive for which they were playing. They then performed the motor task, with the same success criteria as during the unincentivized phase. At the end of the experiment, a single trial was selected at random and participants were paid based on their performance on that trial.

To summarize, our task had several important features: 1) During the incentivized performance phase we did not display cursor position to participants so they could not simply target the necessary effort level. Instead they exerted effort in accordance with what they remembered the target effort level to be, and since they were not able to see the target, any extra exertion that they produced captured implicit incentive motivational spill-over into motor performance. 2) We parametrically modulated incentive to provide a finer grained assessment of how performance varies with incentive, unlike previous investigations of motor cortical influences on instrumental performance, which used appetitive and aversive conditioned stimuli (in extinction) and were not designed to examine parametric effects of rewards (Chiu et al., 2014; Freeman et al., 2014; Freeman and Aron, 2016). Notably, the previous TMS study that did parametrically vary incentive was designed to study decision-making and not incentive motivated instrumental performance (Klein-Flügge and Bestmann, 2012). 3) To evaluate the influence of subjective preferences on incentive motivation we had participants perform a separate prospect theory task that provided a precise measurement of subjective preferences for incentive (i.e. loss aversion, risk aversion) (Sokol-Hessner et al., 2009; Chib et al., 2012, 2014). This task generated measures of subjective preferences for reward that were independent of the incentive motivation task, which allowed an unbiased means to examine relationships between sensitivity to incentive, incentive motivated behavior, and motor cortical excitability.

### Subjective Reward Preference Task (Measurement of Loss Aversion and Risk Aversion)

Participants received an initial endowment of $25 in cash (this amount was separate from their show-up fee and earnings/endowment from the testing phase) and were told that, at the end of the experiment, one trial would be selected randomly and a payment made according to their actual decision during the experiment. Participants were told that their $25 endowment was given to them so that they could pay any eventual losses at the end of the experiment. Any net amount from the endowment that remained after subtracting a loss was theirs to keep, and similarly any eventual gain earned in the experiment was added to the initial endowment. During the experiment, participants made choices among 140 different pairs of monetary gambles. Each pair contained a certain option involving a payout with 100% probability, S, and a risky option involving gain, G, and loss, L, with equal probability. Participants had 4 s to make a choice. Specifics of the gambles used can be found in previous studies (Sokol-Hessner et al., 2009; Frydman et al., 2011). These gambles and task have been used in recent studies to obtain measures of individuals’ loss aversion parameters.

### Data Analysis

#### Behavioral performance analysis

Our main behavioral measure of performance was the mean effort exerted on each trial, defined as mean force between an exertion threshold (i.e., the first recording above 10% of MVC) and the end of the trial. We excluded trials if detected reaction time intersected with MEP onset. We used a general linear model with the magnitude of potential gain and loss *x* and valence *v* as independent variables, and performance as the dependent variable.

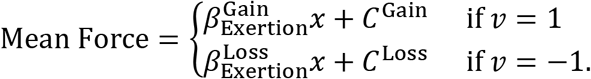

The regression coefficients 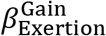 and 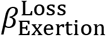 represent a participant’s sensitivity in performance to increasing potential gains and losses — larger *β* parameters correspond to a participant having greater increases in performance as a function of increasing incentives. The parameters C^Gain^ and C^Loss^ capture the performance off-set associated with each valence condition.

### Motor cortical excitability analysis

We assessed cortical excitability by measuring the peak-to-peak amplitudes (in mV) of the motor evoke potential from the FDI muscle on all stimulation trials. This measure was defined as the MEP. In a similar fashion to the behavioral analysis, we used a general linear model to examine the sensitivity of motor cortical excitability to reward, at 50ms and 150ms following the ‘Go’ cue. In this model the magnitude of potential gain and loss *x*, valence *v*, and stimulation time *t* were independent variables; and MEP was the dependent variable. To account for differences in MEP between participants, we mean corrected participants’ MEP measures within session.

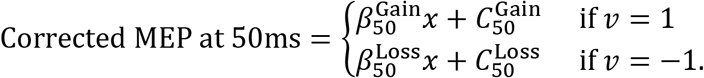

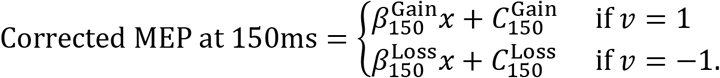

The coefficient terms 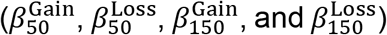 represent an individual’s motor cortical sensitivity to incentive at different time points prior to movement. The intercept terms (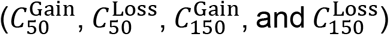 capture the MEP offset associated with each valence condition.

### Subjective reward preferences analysis

We fit prospect theory-inspired models of the non-linear processes underlying subjective valuation of reward to participant’s choice data from the subjective reward preference task, using a hierarchical Bayesian approach. This model was identical to that used previously (Sokol-Hessner et al., 2009; Chib et al., 2012, 2014; Ahn et al., 2017). We expressed participants’ utility function *u*(*x*) for monetary values *x* as follows:

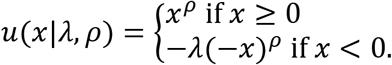

This formulation is used to compute the utilities of the risky and certain alternative. The model’s parameters quantify loss aversion (*λ*, the relative multiplicative weight placed on losses compared with gains), risk attitudes (ρ, feelings about chance, or diminishing marginal sensitivity to value). Assuming participants combine probabilities and utilities linearly, the expected utility of a mixed gamble can be written as *U*(G, L,S|*λ,ρ*) = .5(G^*ρ*^ – *λ*L^*ρ*^) – S^*ρ*^ where G and L are the respective gain and loss of a presented risky option and S is a fixed alternative choice. The probability that a participant chooses to make a gamble is given by the softmax function:

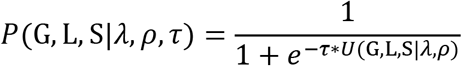

where τ is a temperature parameter representing the stochasticity of a participant’s choice (τ = 0 means choices are random). This model was fit to the choice data using standard Markov-Chain Monte Carlo sampling methods in rStan (v2.2.0; (Stan Development Team, 2017)) as implemented in R (v3.0.2; (RDevelopment CORE TEAM, 2008)). All analyses of loss aversion used log(*λ*); the logarithm is commonly used because lambda is positively skewed.

### Mediation analysis

Mediation analysis is a specific case of structural equation modeling that refers to a situation that includes three or more variables, such that there is a causal process between all three variables (Judd and Kenny, 1981). In a mediation relationship, there is a direct effect between an independent variable and a dependent variable. There are also indirect effects between an independent variable and a mediator variable and between a mediator variable and a dependent variable. This formulation allows for a test of the strength of the direct effect between the independent and dependent variables, accounting for connections via a mediating variable. A measure of the direct effect (after controlling for the mediator) can be obtained using a series of regressions for all of the causal pathways and estimating a change in the direct effect.

We performed a mediation analysis of our data to test the possibility that the relationships between subjective preferences for reward (instantiated by loss aversion), and task performance were mediated through motor cortical excitability. For these analyses, we performed between-participant regressions with variables for participants’ behavioral loss aversion, the difference in motor cortical sensitivity to prospective gain between stimulation at 50ms and 150ms 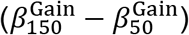), and the performance sensitivity to increasing potential gain 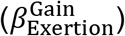). Our main mediation hypothesis was that 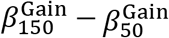 mediates the relationship between *λ* and performance. To rule out model misspecification, we also tested control models in which the causal structure of our experiment was preserved (i.e., motor excitability preceded performance), and alternative relationships were modeled. This included a model in which *λ* mediated the relationship between 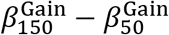 and 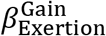, and another model in which 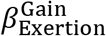 mediates the relationship between 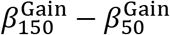 and *λ*. We used bootstrapping (a nonparametric sampling procedure) to test whether the specified mediator significantly mediated the relation between the independent and dependent variables (Preacher and Hayes, 2004).

## RESULTS

To test our hypotheses, we developed a task in which participants were instructed to exert pinch grip beyond a predetermined threshold in order to win or avoid losing monetary incentives ranging from $0 to $20. We stimulated participants’ motor cortex with TMS at two timepoints, between the presentation of incentive and movement onset, in order to examine how motor cortical sensitivity to incentive was related to incentive motivation. Participants also performed a separate decision-making task after performing the motor task, in which they made choices over prospective monetary gains and losses. This task allowed us to obtain computational parameters that described each participants’ subjective preferences for incentive (i.e., loss aversion and risk aversion).

To foreshadow the results, we found that participants exhibited increasing behavioral performance for increasing incentives, and that these increases in performance were related to motor cortical sensitivity to incentive in the time period between incentive presentation and movement. Both performance and motor cortical sensitivity to incentive were related to measures of participants’ loss aversion, such that those individuals that were more loss averse (i.e., had a greater sensitivity to incentive) exhibited larger behavioral and motor cortical sensitivity to incentive. A formal mediation analysis revealed that motor cortical sensitivity to incentive mediated the relationship between subjective preferences for incentive and performance.

## Behavioral Performance

As expected, prospective gains and losses led to increases in participants’ percent success when comparing $0 trials to $10 and $20 trials (Figure 2A; Wilcoxon signed rank paired test to account for skewed distribution at $10 and $20, Gain: *Z* = 111.50; *p* = 0.05; Loss: *Z* = 135; *p* = 0.0018). We also observed robust relationships between participants’ mean exertion as a function of incentive. We found that participants also exhibited increasing mean exertion, with increasing incentives, in both the gain and loss conditions (Figure 2B; hierarchical linear regression, Gain; *β* = 0.018, *t*(104) = 3.8, *p* = 2.6 * 10^-4^; Loss; *β* = 0.025,*t*(104) = 7.0, *p* = 2.9 * 10^-10^). Together, these results illustrate that increasing incentives serve to increase behavioral performance in both the gain and loss domain.

**Figure 2.**
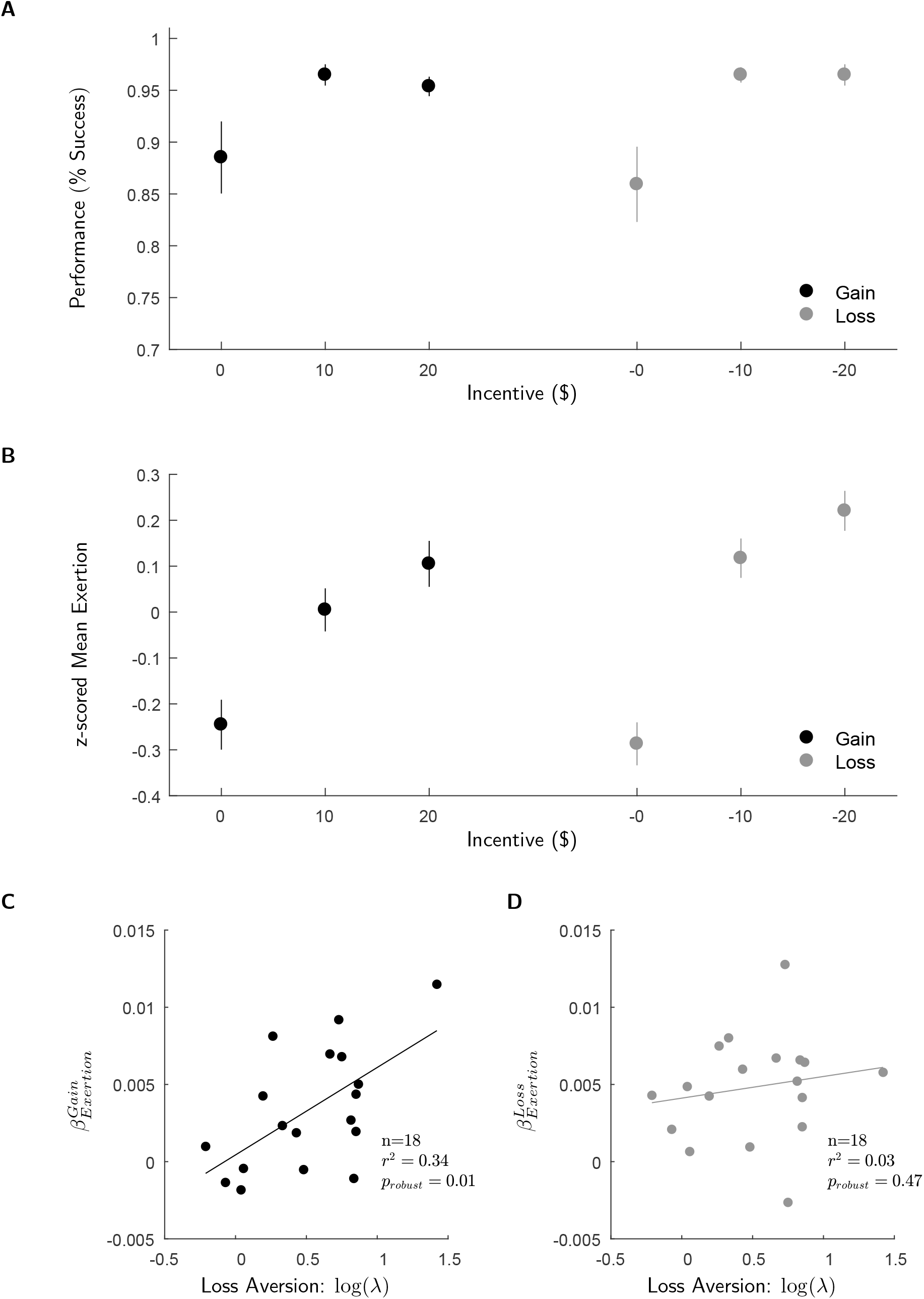
Behavioral results. **A**) Participants exhibited increasing performance (% success) for increasing prospective gains and losses. **B**) Participants exerted more pinch force (mean effort exertion) for increasing prospective gains and losses. Mean exertion was z-scored to control for inter-participant variability in performance. Plots of the correlation between participants’ behavioral sensitivity to prospective (**C**) gains and (**D**) losses (i.e., slope of the relationship between un-normalized mean exertion and incentive) and loss aversion. Error bars denote SEM.

We next examined the relationship between participants’ behavioral sensitivity to increasing prospective gains and losses in the incentive motivation task, and an independent measure of participants’ sensitivity to incentive obtained from a separate decision-making task. We reasoned that those individuals that found incentives to be more subjectively valuable (i.e., have a higher loss aversion) would have increased behavioral sensitivity to incentive. We found a significant relationship between participant specific loss aversion and behavioral sensitivity in the gain domain (Figure 2C; robust regression, *t* (16) = 2.8, *p* = 0.013), however we failed to find a significant relationship between these measures in the loss domain (Figure 2D; robust regression, *t*(16) = 0.74, *p* = 0.47). This suggests that, for prospective gains, processing of the subjective value of incentive serves to motivate behavioral performance in the incentive motivation task. These results align with our previous work which found that loss aversion was predictive of increases in performance for incentives in the range tested in this experiment (Chib et al., 2012, 2014). In those previous studies, we found that worries about loss (instantiated by loss aversion) served to motivate performance for both prospective gains and losses.

Loss aversion represents a tendency to value losses greater than equal magnitude gains. Risk aversion, on the other hand, is a more general aversion to increased variance in potential gains or losses. To ensure a loss aversion-based hypothesis, and not a general aversion to risk was responsible for our findings, we also examined the relationship between risk aversion and behavioral sensitivity in the gain and loss domains. We did not find a significant correlation between behavioral sensitivity to incentive and risk preferences (robust regression, Gain: *t*(16) = –0.78, *p* = 0.45; Loss: *t*(16) = –1.3, *p* = 0.21). This provides further evidence that behavioral sensitivity to prospective gains is solely dependent on reward subjectivity characterized by a measure of loss aversion.

## Motor Cortical Excitability in Response to Incentive

We sought to identify how motor cortical sensitivity to incentive, in the context of the incentive motivation task, was related to subjective preferences for incentive. To this end we explored parameter estimates from our general linear model of motor cortical sensitivity to incentive, separated by participants’ behavioral loss aversion (participant specific medial split) (Figure 3A). These parameter estimates capture the slope of the relationship between motor cortical excitability and incentive level. Larger parameter estimates correspond to a more pronounced change in motor cortical excitability in response to increasing incentive.

**Figure 3.**
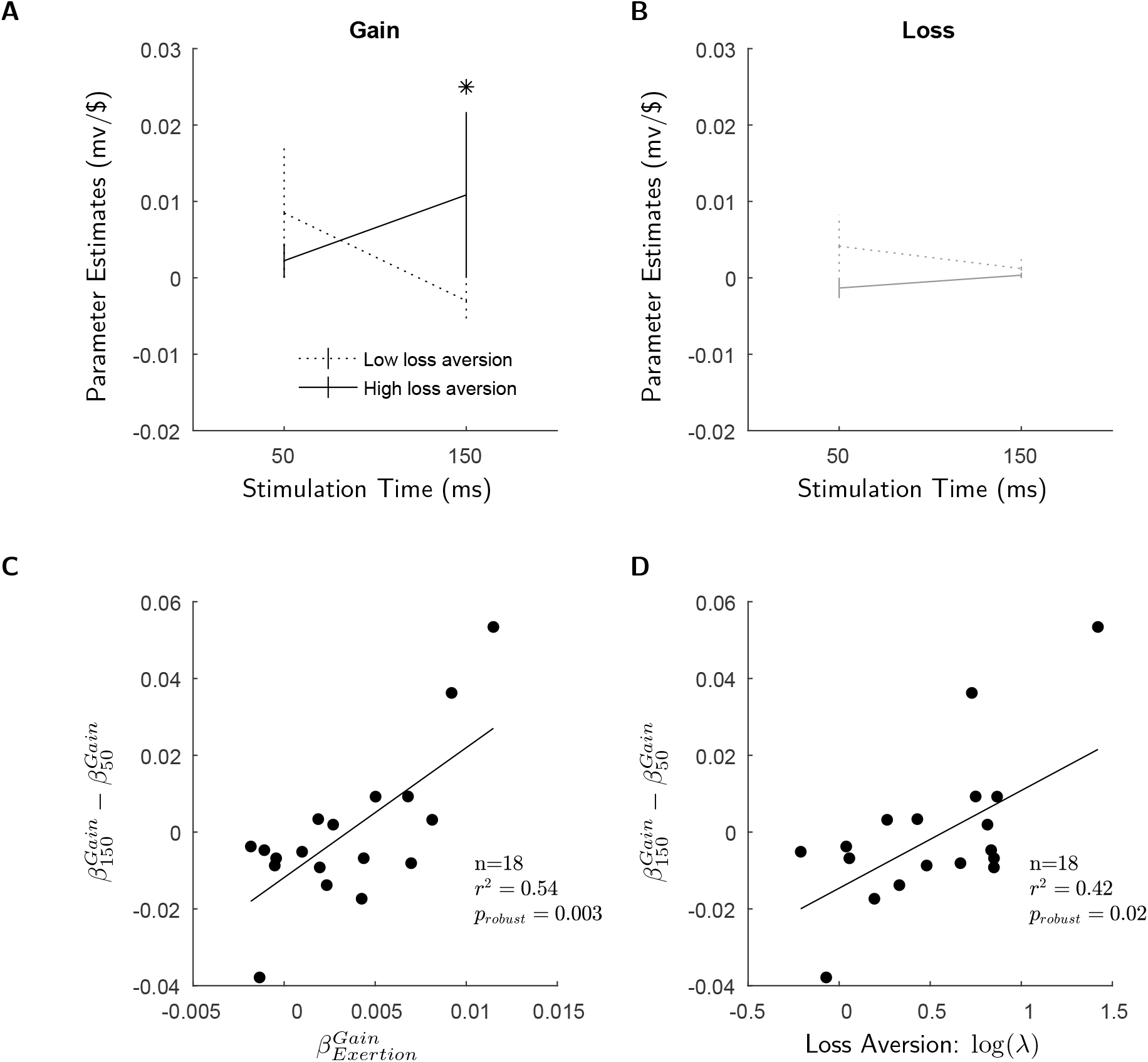
Motor cortical excitability in response to incentive. Shown are fixed effects parameter estimates from hierarchical linear regression models predicting motor cortical sensitivity to incentive at the different TMS time points. Positive parameter estimates indicate increasing motor excitability with increasing incentive, negative estimates indicate decreasing motor excitability with increasing incentive, and zero estimates indicate no modulation of motor cortical excitability with incentive. **A, B**) We separated trials based on prospective gain and loss, and grouped participants by the extent of their loss aversion (median split). In the gain domain, we found that those participants that were more loss averse had greater increases in motor cortical excitability in response to incentive, closer to movement imperative (150ms). We failed to find significant modulation of motor cortical excitably for prospective loss. The significance levels shown are for planned comparisons between conditions (*p < 0.05). Error bars denote SEM. Plots of the correlations between difference in motor cortical sensitivity to incentive between the 150ms and 50ms stimulation conditions, in the gain domain, and (**C**) behavioral sensitivity to incentive (i.e., slope of the relationship between un-normalized mean exertion and incentive) and (**D**) behavioral loss aversion.

In the gain domain, we found a significant interaction between stimulation time and loss aversion, indicating that individuals with higher loss aversion had an increased motor cortical sensitivity to incentive, closer to movement initiation (Figure 3A; ANOVA, *F*(1,32) = 6.1, *p* = 0.019). Moreover, we found that this effect was driven by individuals with higher loss aversion having an increased motor cortical sensitivity at the 150ms stimulation time point (Figure 3A, post hoc Welch’s t-test, *t*(10.37) = 2.54, *p* = 0.030). In the loss domain, we failed to find a significant interaction between changes in MEP sensitivity between the 50 and 150 stimulation time points and behavioral loss aversion (Figure 3B; ANOVA, *F*(1,32) = 1.1, *p* = 0.3), or modulation of motor cortical activity in response to increasing prospective losses.

Given that we failed to find significant effects in the loss domain, we reasoned that our experimental paradigm was not sensitive to potential changes in motor excitability in response to prospective losses. It should be noted that this null result is consistent with a previous study of motor cortical responses to aversive stimuli which failed to find a significant change in motor evoked potentials, relative to baseline, when individuals were presented an aversive conditioned stimulus paired with an instrumental response (Chiu et al., 2014). With this null result in the loss domain in mind, we focused the remainder of our motor cortical analyses on trials for prospective gain.

To further examine the temporal dynamics of motor cortical sensitivity to incentive over the continuum of loss aversion, we performed a between participant regression of loss aversion and difference in sensitivity to incentive between the 50ms and 150ms time points 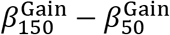. The difference between these metrics is an indication of the stability of motor cortical excitably to incentive over time. The greater the difference between these parameter estimates, the more positively correlated to incentive an individual’s motor cortical excitably is closer to movement onset. We found that those individuals that were more sensitive to incentive when comparing 150ms and 50ms time points, exhibited increased incentive motivated performance (Figure 3C; robust regression; *β* = 3.4, *t*(16) = 3.5, *p* = 0.0027). We also performed a regression between participant-specific loss aversion and sensitivity to incentive between the 50ms and 150ms time points and found that individuals with higher loss aversion exhibited increased changes in motor cortical sensitivity closer to movement imperative (Figure 3D; robust regression; *β* = 0.026, *t*(16) = 2.7, *p* = 0.016).

In keeping with our incentive motivation hypotheses of motor cortical activity, these relationships suggest that in the gain domain, subjective preferences for incentive (instantiated by an individuals’ loss aversion) could serve to amplify motor cortical sensitivity to incentive and energize motor performance.

We performed a series of analyses to ensure that the TMS incentive effects that we observed were not simply the byproduct of confounds between stimulation timing and movement execution. Pre-movement motor cortical stimulation is known to elicit movement quickening, in which stimulations delivered closer to movement imperative decrease reaction time. To ensure that our TMS incentive effects were not simply the byproduct of a quickening response, we examined the relationship between reaction time sensitivity to incentive (i.e., the regression coefficient between reaction time and incentive) and motor cortical sensitivity to incentive, at each stimulation time point, and using our measure of motor cortical sensitivity to incentive 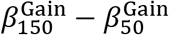. We failed to find significant correlations between these measures suggesting that our effects were not simply the results of TMS quickening movements as a function of incentive (Figure 4A, robust regression; *β* = −2.1, *t*(16) = −1.2, *p* = 0.26).

**Figure 4.**
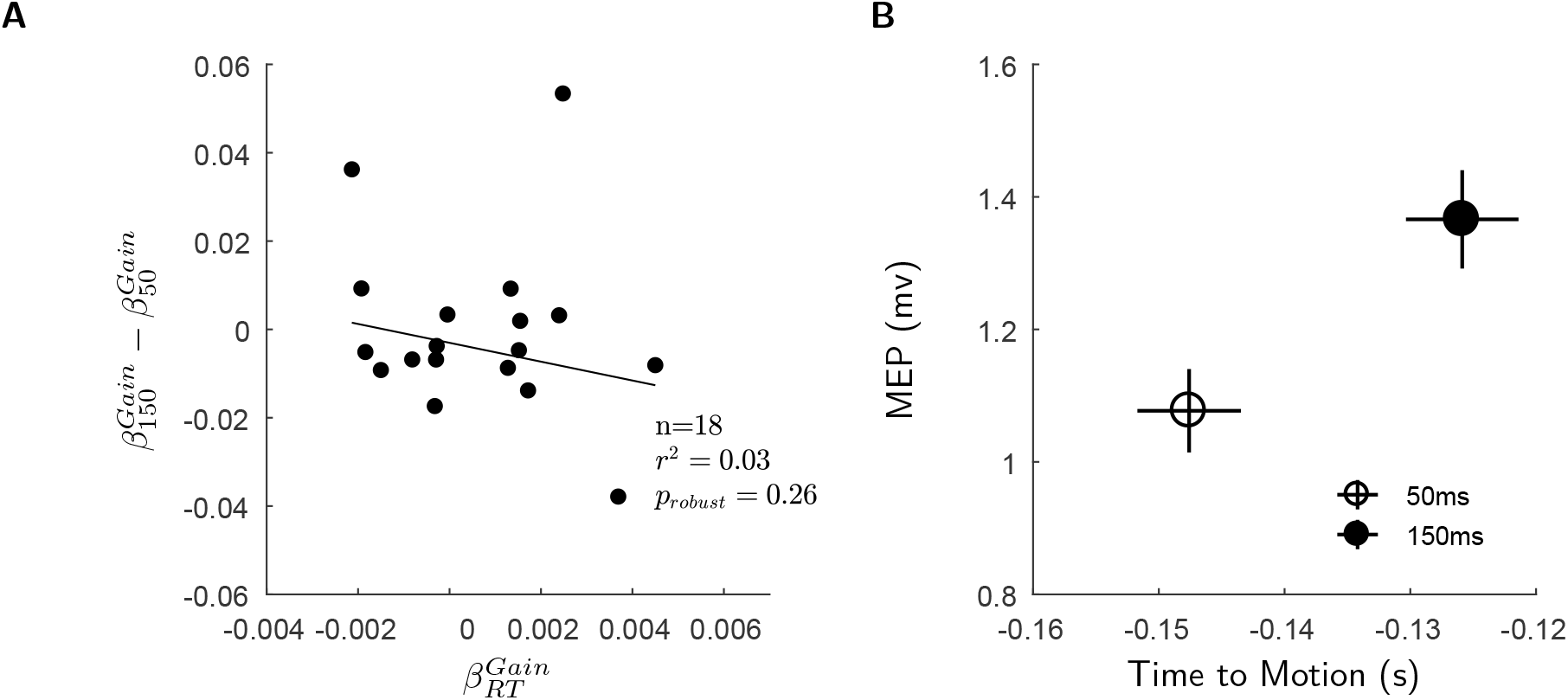
Control TMS analyses. **A**) We did not to find a significant relationship between difference in motor cortical sensitivity to incentive between 50ms and 150ms and reaction time sensitivity to incentive. **B**) Motor evoked potentials were segregated in intensity and time when aligning them to EMG detected movement onset, rather than the ‘Go’ cue.

Another possible confounding factor in our motor cortical data could be that participants initiate their movements based on the auditory cue of TMS pulses, rather than the ‘Go’ cue. This would result in no segregation between motor cortical activity between the 50ms and 150ms stimulation conditions, making it difficult to distinguish the temporal features of motor cortical sensitivity to incentive. To determine if our data was confounded in this way, we evaluated motor cortical excitability using a model in which trials were separated based on the eventual time of movement onset (as identified from participants EMG data using AGLRStep (Staude et al., 2001)), rather than presentation of the ‘Go’ cue (as in our main experimental results). We found that, although there was some quickening as a result of TMS (i.e., MEPs were not separated by a full 100ms), motor evoked potentials occurred at significantly different time points relative to movement onset (Figure 4B; paired t-test; *t*(17) = 15.0, *p* = 2.0 * 10^-11^). Moreover, we found that motor evoked potentials were larger in the 150ms stimulation condition compared to the 50ms condition (Figure 4B; paired t-test; *t*(17) = 4.7, *p* = 2.3 * 10^-4^), consistent with previous studies that have shown increasing motor cortical excitability approaching movement imperative (Chen and Hallett, 1999).

## Causal Influences of Loss Aversion and Motor Cortical Excitability on Incentive Motivation

Because loss aversion and behavioral sensitivity to incentive are correlated, and both of these variables are correlated with the temporal evolution of motor cortical sensitivity to incentive (Figure 5A), we investigated the hypothesis that motor cortical sensitivity to incentive has a causal influence on loss aversion-related incentive motivated exertion. To test this hypothesis, we used mediation analysis, a form of linear modeling in which correlations observed in the data are explained by assuming that a specific set of causal influences exist among the variables (Judd and Kenny 1981). This analysis alone does not establish causality but identifies if a causal hypothesis is best fit for the data. We fit a model to the data that followed the logical progression of our experimental paradigm. In this model we assumed that behavioral loss aversion influenced incentive motivated exertion and that the temporal dynamics of motor cortical sensitivity to incentive (the mediating variable) influenced incentive motivated exertion.

**Figure 5.**
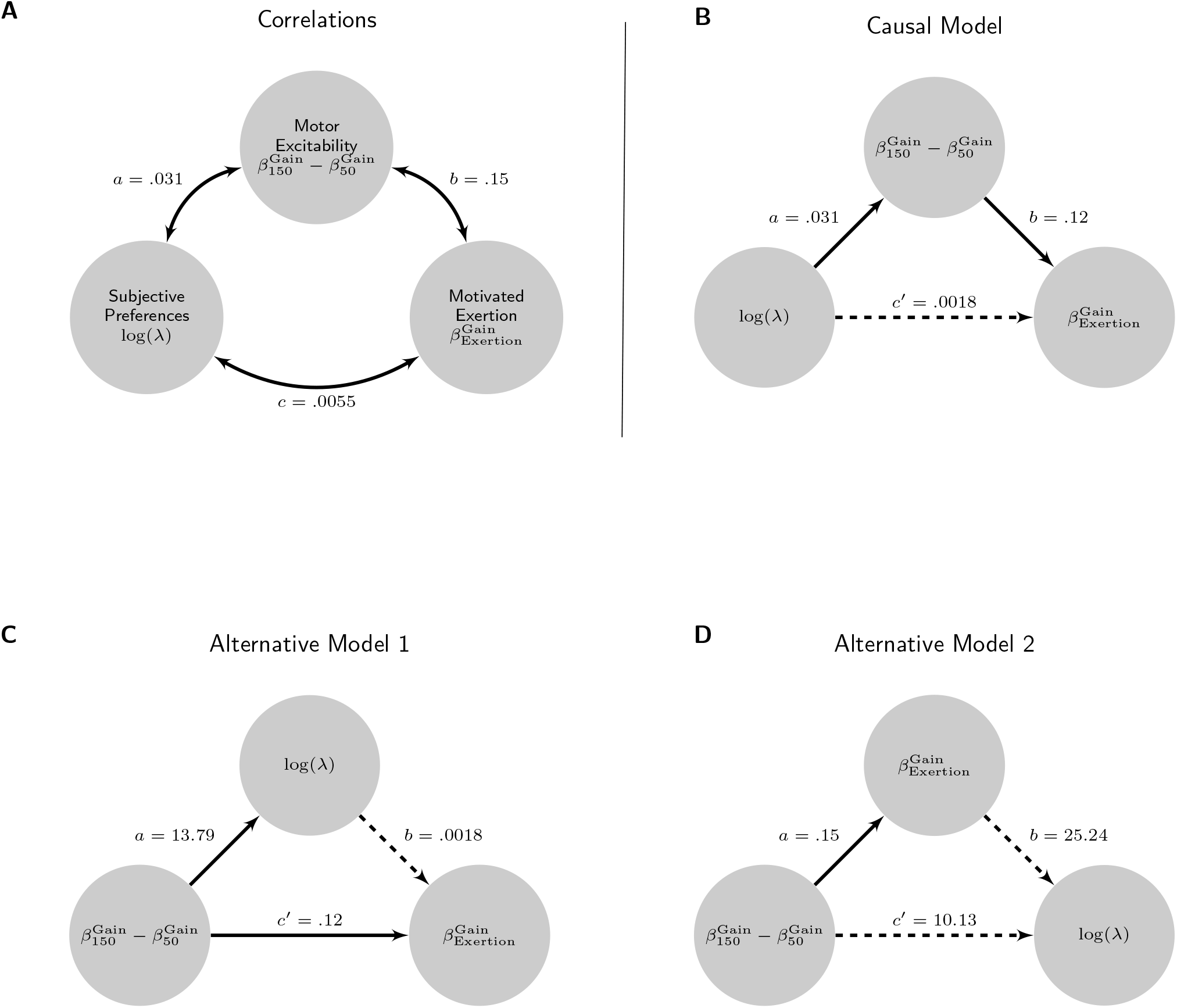
Mediation analyses. **A**) The three variables assessed using mediation analysis: behavioral loss aversion log(*λ*) difference in motor cortical sensitivity to incentive between the 50ms and 150ms TMS time points 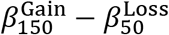, and behavioral sensitivity to incentive 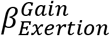. The numbers next to the double-headed arrows are coefficients of correlations between the variables. Regression analyses (illustrated in Figures 2C,3C,3D) established correlations between participants behavioral loss aversion, differences in motor cortical sensitivity to incentive, and behavioral sensitivity to incentive. **B**) The causal model illustrates the mediation analysis, and the alternative models (**C, D**) illustrate control models that were tested to rule out model misspecification. Solid arrows represent significant relationships between variables, dashed arrows are not significant.

In our causal model (Figure 5B), behavioral loss aversion had a significant effect on the difference in motor cortical sensitivity to incentive between the 50ms and 150ms time points (linear regression, *β* = 0.031, *t*(16) = 3.4, *p* = 0.0034). When behavioral loss aversion and this measure of motor cortical sensitivity to incentive were simultaneously modeled as predictors of performance, loss aversion no longer significantly predicted performance (linear regression, *β* = 0.0018, *t*(15) = 0.85, *p* = 0.41), whereas motor cortical sensitivity to incentive remained significant in the model (linear regression, *β* = 0.12, *t*(15) = 2.7, *p* = 0.017). This reduction in the direct relationship between loss aversion and incentive motivation was significant (95% confidence interval, 2.2 * 10^−4^ to .0080; *p* < 0.05, as tested by a bootstrapping procedure based on 10000 resamples). This model provides causal support for the idea that manifestations of subjective preferences for incentive motivate incentivized performance through the influence of the temporal dynamics of motor cortical sensitivity on motor performance. Our alternative models (Figure 5C) ruled out model misspecification and did not find significant mediation effects by loss aversion (95% confidence interval, –0.038 to .0090; *p* ≮ 0.05) or performance (95% confidence interval, −5.2 to 13.1; *p* ≮ 0.05).

## DISCUSSION

In this study we show that incentive motivated performance emerges from the temporal dynamics of motor cortical sensitivity to incentive, and that this signature of motor cortical activity reflects an individual’s subjective preferences for incentive and eventual behavioral performance. Our neural findings are consistent with previous evidence showing that appetitive stimuli serve to increase motor cortical excitability (Chiu et al., 2014; Freeman et al., 2014) and that the dynamics of motor cortical excitability is sensitive to the value of options presented during simple choice (Klein-Flügge and Bestmann, 2012). However, as previous studies either investigated instrumental responding or value-based choice in separate paradigms; they did not examine the relationship between the temporal dynamics of motor cortical sensitivity to incentive, subjective preferences for incentive, and eventual motor performance. Our results go beyond these studies by separately characterizing the temporal dynamics of motor cortical sensitivity to incentive and subjective preferences for incentive, and further, modeling the causal relationship between these independent measures and behavioral performance. In so doing, we demonstrate a mechanism by which motor cortical activity mediates the relationship between subjective preferences for incentive and incentive motivated performance. These results suggest that an individual’s subjective preferences for incentive modulate the vigor of the motor system to drive incentive motivated performance.

We previously used functional imaging to show that when performing an instrumental motor task for incentive, prospective incentives are first encoded as a potential gain and subsequently, during the task itself, individuals encode the potential loss that would arise from failure (Chib et al., 2012, 2014). This reframed loss encoding served to motivate behavioral performance — those individuals that were more loss averse had a greater behavioral sensitivity to incentive, such that they reached peak performance at lower incentive levels. Moreover, we found that activity in the ventral striatum, a region of the brain thought to serve as the interface between motivation and motor performance (Mogenson et al., 1980; Bray et al., 2008; Talmi et al., 2008), was predictive of both performance and loss aversion. Consistent with these results, here we found that behavioral sensitivity to incentive in the gain domain was related to an individuals’ loss aversion. Those individuals that were more loss averse had increased behavioral sensitivity to incentive, suggesting they were more motivated for increasing incentives. The temporal dynamics of motor cortical sensitivity to incentive also reflected an individual’s behavioral loss aversion – those individuals that were more loss averse showed an increasing motor cortical sensitivity to incentive closer to movement imperative. These new TMS results take our previous reframing interpretation further and show that motivational constructs (i.e. loss aversion), known to be encoded by reward regions of the brain, transfer to motor areas (as reflected by motor cortical excitability changes), giving rise to motivated behavioral performance.

The present results provide important new insights into how incentive motivational processing influences motor cortical activity to give rise to performance. One possible mechanistic account of our findings relates to the role of the ventral striatum as a limbic motor interface, mediating interactions between systems for Pavlovian valuation and motoric instrumental responding (Mogenson et al., 1980; Alexander and Crutcher, 1990; Cardinal et al., 2002; Balleine and Ostlund, 2007). Whereas previous literature has focused on the role of the ventral striatum in mediating the effect of reward-predicting cues in increasing or enhancing instrumental performance for reward, less is known about how such reward processing influences activity in motor cortex to give rise to behavioral performance. An elegant set of studies used a Pavlovian instrumental transfer paradigm to study such effects, and showed that appetitive cues served to increase motor cortical excitability during instrumental responding (Chiu et al., 2014; Freeman et al., 2014). In our experiment, it is possible that during motor performance the prospect of reward (and loss-aversion induced motivation) elicits participants’ Pavlovian conditioned responses. These responses could include motor approach and engagement of attentional or orienting mechanisms towards task performance. Such ventral striatal encoding of Pavlovian responses could energize the motor cortical commands necessary for successful execution of instrumental responses, and this motor energization could manifest in the motor cortical sensitivities to incentive that we observe in our data. Accordingly, there are strong direct and indirect connections between ventral striatal regions known to encode such Pavlovian and reward values and motor cortex (Mogenson et al., 1980; Haber and Knutson, 2010).

Further supporting these ideas about the motor cortex, was a mediation analysis showing that motor cortical sensitivity to incentive mediated the effects of behavioral loss aversion on performance. This mediation suggests that the motor cortex is not merely indirectly correlated with performance through its relationship with loss aversion, but instead plays a critical role in moderating incentive motivated behavioral performance itself. This provides a mechanistic account of how the motor cortex influences motivated motor performance via its reflection of subjective preferences and incentive value.

It is important to note that although we found a significant modulation of behavioral performance for increasing prospective loss, we failed to find such an effect in motor cortical excitability responses. Notably, a previous study that examined how aversive conditioned stimuli influenced motor cortical excitability, during instrumental responding, also failed to find a modulation of motor cortical excitability by aversive stimuli (Chiu et al., 2014). One interpretation of these null results is that distinct neural circuits could process the effects of appetitive and aversive stimuli on motivated motor performance (Pessiglione and Delgado, 2015). Indeed, distinct amygdala nuclei have been shown to encode appetitive (basolateral amygdala) (Holland et al., 2002) and aversive (central nuclei) (Petrovich et al., 2009) stimuli during motivated behavior. These amygdala nuclei are essential components in the circuits that mediate Pavlovian instrumental transfer and have different circuit pathways that connect to ventral striatum to influence motivated performance (Cador et al., 1989; Corbit et al., 2001; Lingawi and Balleine, 2012). However, it is not known if these pathways also have different connections to the motor cortex. It is possible that such differential TMS effects could be the result of such distinct pathways for appetitive and aversive stimuli. Resolving this possibility is beyond the design of the current study and could be achieved using functional neuroimaging techniques, combined with noninvasive brain stimulation, to examine how motor cortical excitability is related to amygdala and ventral striatal function in the context of motor performance for prospective gains and losses.

Integrating behavioral analysis of motivated performance, modeling of subjective preferences for incentive, and motor cortical physiology; we provide evidence that the motor cortex is sensitive to the subjective value of incentive. Our work outlines a mechanism by which the subjective value of reward serves to invigorate motor cortical excitability, leading to incentive motivated performance. Far from simply being a reflection of motor output, it appears that motor cortical physiology integrates cognitive mechanisms related to reward valuation. These results suggest that incentive motivated performance is the reflection of an interaction between reward valuation and motor cortical excitability.

## Acknowledgements

This work was supported by the Eunice Kennedy Shriver National Institute Of Child Health & Human Development of the National Institutes of Health under Award Number K12HD073945 to V.S.C. J.G. was supported by the National Defense Science and Engineering Graduate Fellowship.

